# Post-intervention Epidemiology of STH in Bangladesh: data to sustain the gains

**DOI:** 10.1101/2020.07.17.208355

**Authors:** Sanjaya Dhakal, Mohammad Jahirul Karim, Abdullah Al Kawsar, Jasmine Irish, Mujibur Rahman, Cara Tupps, Ashraful Kabir, Rubina Imtiaz

## Abstract

**Introduction:** In 2008, Bangladesh initiated Preventive Chemotherapy (PCT) for school-age children (SAC) through bi-annual school-based mass drug administration (MDA) to control Soil-Transmitted Helminth (STH) infections. In 2016, the Ministry of Health and Family Welfare’s Program on Lymphatic Filariasis Elimination and STH (ELFSTH) initiated district-level community impact assessments with Children Without Worms (CWW) using standardized, population-based sampling to measure the post-intervention STH burden across all ages (≥ 1 yr) for the three STH species.

**Methods & Principal Findings:** The Integrated Community-based Survey for Program Monitoring (ICSPM) was developed by CWW and was used to survey 12 districts in Bangladesh from 2017 – 2020. We combined the individual demographic and parasite-specific characteristics from 10 districts and linked them with the laboratory data for collective analysis. Our analysis identified district-specific epidemiologic findings, important for program decisions.

Of the 17,874 enrolled individuals, 10,824 (61%) provided stool samples. Overall, the prevalence of any STH species was substantially reduced to 14% from 79.8% in 2005. The impact was similar across all ages. STH prevalence was below 10% in 10 districts collectively, but remained high in 4 districts, despite their high reported PCT coverage in previous years. Among all, Bhola district was unique because it was the only district with high Trichuris prevalence.

**Conclusion:** Bangladesh successfully lowered STH prevalence across all ages despite targeting SAC only. Data from the survey indicate significant number of adults and pre-school age children (PSAC) were self-deworming with purchased pills. This may account for the flat impact curve across all ages. Overall prevalence varied across surveyed districts, with persistent high transmission in the northeastern districts and a district in the central flood zone, indicating possible service and ecological factors. Discrepancies in the impact between districts highlight the need for district-level data to evaluate program implementation after consistent high PCT coverage.

**Authors Summary:** Bangladesh government conducted school-based mass drug administration (MDA) for over 10 years to control soil-transmitted helminth (STH) infections. School-based evaluations of MDA indicate a reduction in STH burden among school-aged children (SAC). To further assess the impact on the community, Children Without Worms and the Ministry of Health and Family Welfare’s Program on Lymphatic Filariasis Elimination and STH (ELFSTH) initiated district-level community impact surveys in 12 districts. We share the results from the latter 10 districts here.

Our analysis of 10,824 interviews and stool samples from 10 districts showed an estimated 14% of community members infected with at least one species of STH. This finding is substantially lower than the baseline STH prevalence (79.8%) estimated in 2005. Bangladesh’s successful impact was achieved across all ages despite only treating SAC. Deworming source data showed significant numbers of adults and pre-school age children (PSAC) self-dewormed with locally purchased pills. Prevalence varied across the surveyed districts, with persistent high transmission in the northeastern districts and a district in the central flood zone, indicating possible ecological and service factors contributing to persistent infections. Discrepancies in the impact across districts highlights the need for sub-national level data to evaluate program performance fllowing consistent high intervention.

## Introduction

In 2001, the World Health Organization (WHO) recommended that member states control Soil-Transmitted Helminthiasis (STH) morbidity through preventive chemotherapy (PCT) in endemic regions. The recommended guidance utilizes a school-based platform to target one high-risk group, school-age children (SAC) through mass drug administration (MDA) to achieve at least 75% coverage consistently for five years. Once this is achieved, an impact assessment survey is recommended (1). Like many developing countries, Bangladesh bears a high burden of STH. An estimated STH prevalence of 79.8% (44% of which was moderate-to-high intensity, MHI, of Ascaris) among school-aged Bangladeshi children was reported in 2005 (2). By January 2020, Bangladesh had completed 23 rounds of school-based bi-annual MDA with Mebendazole. Bangladesh receives the largest Mebendazole donation of all endemic countries (approximately 20% of the global donation) and has an excellent supply chain record over the past 5 years (SCF-NTD data: CWW pulled June 2020).

Annual coverage data from Bangladesh indicates consistent coverage of greater than 75% for more than five years before ICSPM surveys began in 2017 (3). Previously, PCT coverage data was used as a proxy to indirectly evaluate the impact of deworming on the STH burden (3, 4). A major limitation of this approach was the inability to assess the true burden of disease in the community at risk because; 1.) MDA targets only SAC (a small proportion of the at-risk population, 2.) the quality of coverage data is not tested, and 3.) targeted parasites have variable sensitivity to the single drug used for MDAs (5, 6). Additionally, PCT coverage data does not include children outside schools and adults. Available evidence indicates that these additional risk populations such as pre-school-age children (PSAC) and adults, particularaly women of reproductive age (WRA) are also at risk of STH infection and share a substantial disease burden (7–9).

Therefore, to better understand the community-level program impact, the ELFSTH Program, the Bangladesh Ministry of Health & Family Welfare (MOHFW) collaborated with Children Without Worms (CWW) to conduct community-level impact assessment surveys called Integrated Community-based Survey for Program Monitoring (ICSPM) from 2017 to 2020. The main objectives of the surveys were:

i. To estimate the statistically valid prevalence of STH infection and prevalence of moderate to high-intensity infection (MHII) in PSAC, SAC and adults (greater than 14 years old), powered to the district level, and
ii. To evaluate potential correlates of STH infection rates including sanitation & hygiene behaviors (household level) and history and source of deworming (individual level).

In this paper, we present the results of concatenated survey data from 10 districts conducted between 2017 and 2020, focusing on parasite- and age-specific prevalence and infection intensity as well as the geographic variation of STH prevalence. This paper also presents how these results are applicable for use by the ELFSTH program towards future program actions, and how this approach can assist other similarly advanced NTD programs around the world.

## Methods

### Study design

ICSPM surveys were conducted in 12 districts, representing 7 out of 8 divisions across Bangladesh, to evaluate the impact of MDA at the community level for each parasite and each risk group; to validate deworming pill intake and pill source within six months before the survey, and to assess the effect of select WASH variables at the household level. Based on age, we defined three risk groups as follows:

- 1 – 4 pre-school age children (PSAC)
- 5 – 14 school-age children (SAC)
- greater than 14 years old (adults)

The districts were selected by the Bangladesh ELFSTH as a good evaluation unit (EU) as the district is the most common administrative unit for implementation decisions where these results could be utilized. Figure 1 shows the years of each survey for the 10 districts.

**Figure 1:**
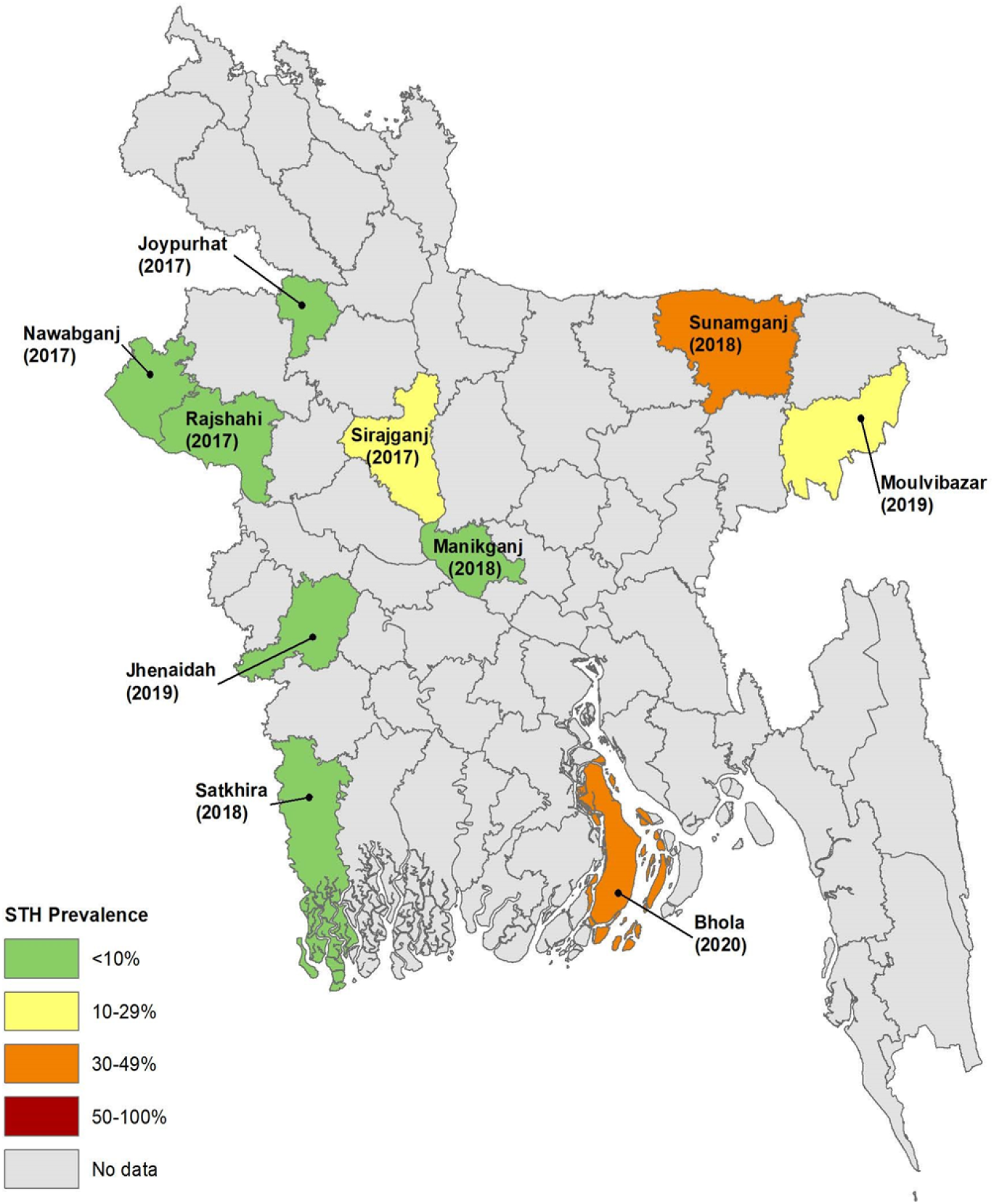
Geographical distribution of ICSPM surveys, year of survey and STH prevalence range.

The ICSPM survey is a community-based, cross-sectional survey based on probability proportional to size sampling (PPSS). The details of the ICSPM survey methodology is available on the CWW website (http://www.childrenwithoutworms.org/). Briefly, the survey design entails a cross-sectional, mixed cluster & random systematic sample methodology and has been previously detailed (10). The ICSPM methodology primarily relies on WHO’s “Assessing the epidemiology of STH during a transmission assessment survey (11). We targeted a sample size of 332 for each risk group, which gave us one-sided 95% confidence for determining if the <10% prevalence, action threshold was achieved. Since the average non-response rate in the first two pilot districts was around 40%, we enrolled 465 individuals in each risk group in subsequent districts to account for this. The sampling interval was based on the proportion of each risk group within the population. The survey team used the Survey Sample Builder (SSB) tool, which was adapted to the ICSPM methodology, an excel program developed by Neglected Tropical Diseases Support Center, The Task Force for Global Health (TFGH) to select clusters and risk groups within the households.

We used the Kato-Katz method to identify and count the eggs of STH parasites following standard WHO methodology using 2-slides per stool specimen. Ten percent of slides were tested blindly by another laboratory scientist for quality control. Three data sets (Household, Individual, and Laboratory) were downloaded from the secure cloud-based data-hosting platform and saved in local computers at CWW, Atlanta. After basic data cleaning, household data were first merged with individual data and later with laboratory data making one linked data file for each district. We recoded and reformatted variables as necessary to align across the districts before combining the individual data files from surveyed districts. Finally, we prepared one analytical data file for this report by stacking 10 individual data files from each of the surveyed districts. According to the 2011 national population census, this analysis represents about 15.5% of the Bangladesh population.

We used SAS version 9.4 (SAS Inc., Cary NC, USA) to manage and analyze the data. We accounted for the cluster sample survey design in all analyses using appropriate SAS procedures. Chi-square (χ²) test was used to assess differences in prevalence between risk groups and p-values ≤ 0.05 were considered significant. Since the survey was powered to detect the prevalence of STH and MHII down to a threshold of <10% at the district level, only upper sided, 95% confidence limits are reported. We also ran some explorative analyses at the sub-district level, which lacked statistical power but provide useful insights for further program actions.

Participation in the survey was voluntary and participants provided verbal consent before the main survey. Bangladesh Medical Research Council approved the survey protocol.

## Results

### Basic characteristics

Of 17,874 enrollees, 11,022 (62.0%) provided stool samples for laboratory examination. In total, 198 (1.6%) records were excluded during the data cleaning process due to;

a. IDs present in only one dataset
b. duplicate IDs with mismatching data across other variables, and
c. data entry errors.

The final dataset had 10,824 records which were used for the analysis presented here (figure 2).

**Figure 2:**
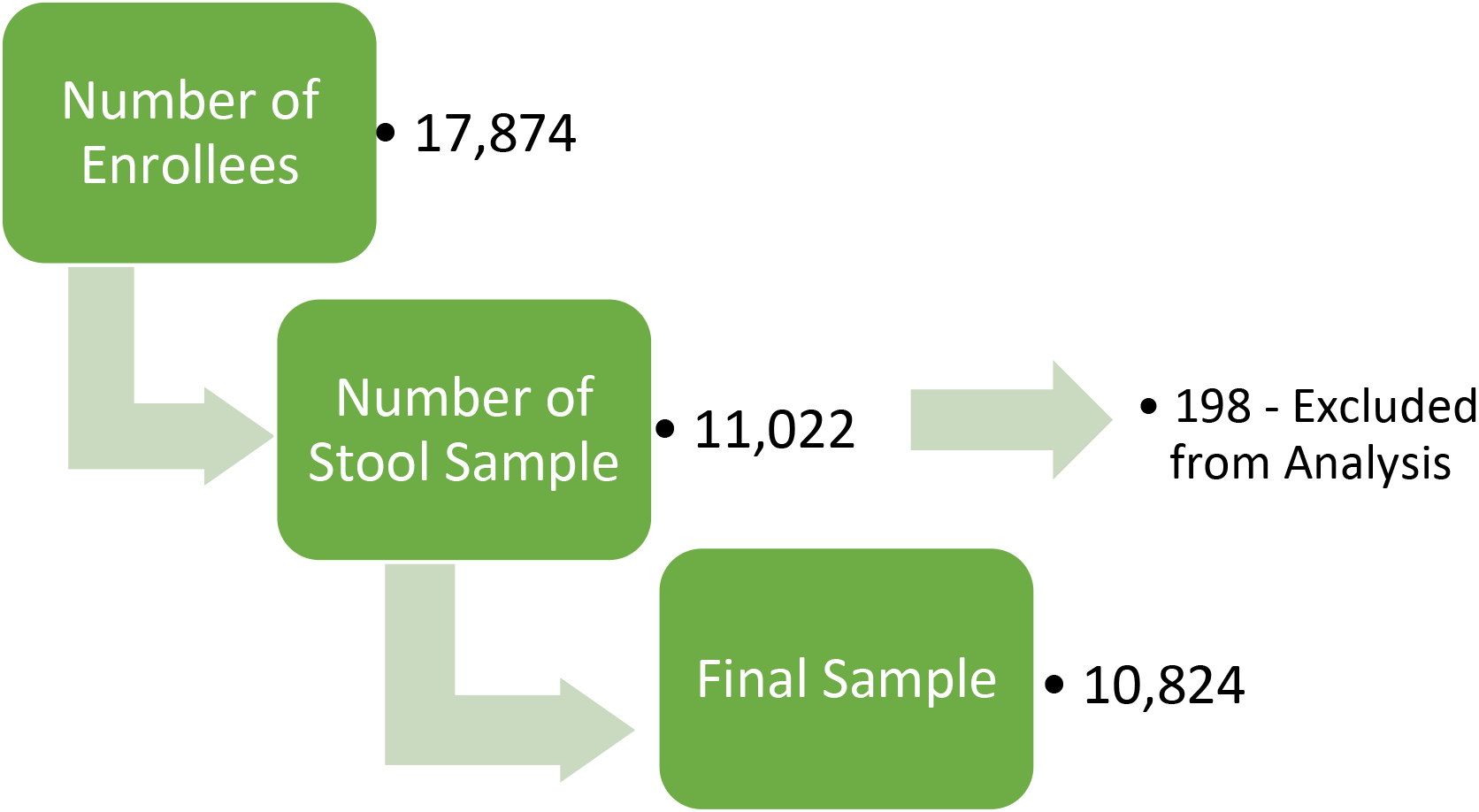
Flow chart of sample selection.

### Prevalence and Intensity of STH Infections

The overall prevalence of any STH infection in 10 districts was 14.0% (figure 3). There was no statistical difference in the prevalence of STH infection across the risk groups. We did not observe statistically different prevalence between females (14.4%) and males (13.4%). Of the three tested parasites, *Ascaris* was the most common (10.5%) followed by *trichuris* (4.4%). The prevalence of hookworm was less than 1% in all risk groups, so the results for hookworm prevalence are not shown. Three districts with the highest STH prevalence were Sunamganj (40.4%), Bhola (36.5%), and Sirajganj (26.9%). In contrast, Satkhira (2.0%), Jhenaidah (2.4%), and Manikganj (3.1%) had the lowest STH prevalence (figure 3).

**Figure 3:**
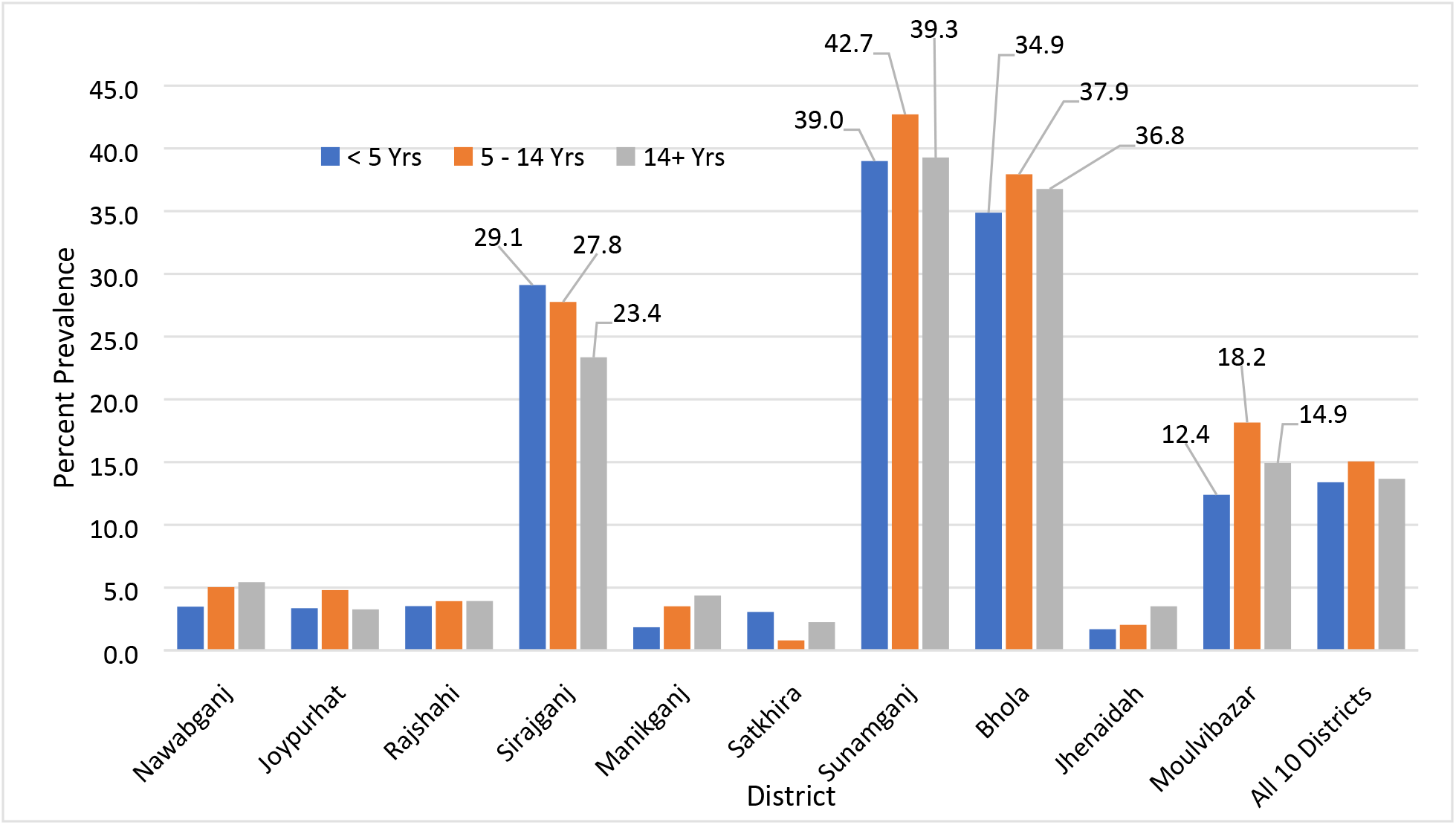
Prevalence of any STH infection by district and age group.

Overall, the intensity of STH MHII for 10 districts was 3.3%. Bhola (10.6%), Sunamganj (10.4%), Sirajganj (7.1%), and Moulvibazar (3.6%) were four districts with MHII above the WHO-recommended threshold of <1%, while the remaining 6 districts had achieved this goal with MHII ranging from 0.0% to 0.2% (table 1). This is a significant achievement for the national program and signifies the achievement of the WHO goal of eliminating STH morbidity in majority districts.

**Table 1:** Intensity of STH morbidity by district

| District | Intensity of STH Morbidity among All (Prevalence %) |  |  | All Ages (%) |
| --- | --- | --- | --- | --- |
|  | PSAC | SAC | Adults |  |
| Sirajganj | 7.0 | 6.6 | 7.8 | 7.1 |
| Sunamganj | 12.2 | 8.6 | 10.2 | 10.4 |
| Bhola | 12.0 | 11.1 | 8.5 | 10.6 |
| Moulvibazar | 3.6 | 4.8 | 2.4 | 3.6 |
| **All 10-districts** | **3.6** | **3.4** | **2.7** | **3.3** |

We further explored three high prevalence (>20%) districts-Sunamganj, Bhola, and Sirajganj- to understand if there were any geographic concentrations of STH infection at the sub-district level. Figure 4 illustrates the STH prevalence by sub-districts in these three high-prevalence districts. The prevalence of STH was higher than 50% in two sub-districts (Dowara Bazar and Dakshin Sunamganj) of Sunamganj and one sub-district (Belkuchi) of Sirajganj district.

**Figure 4:**
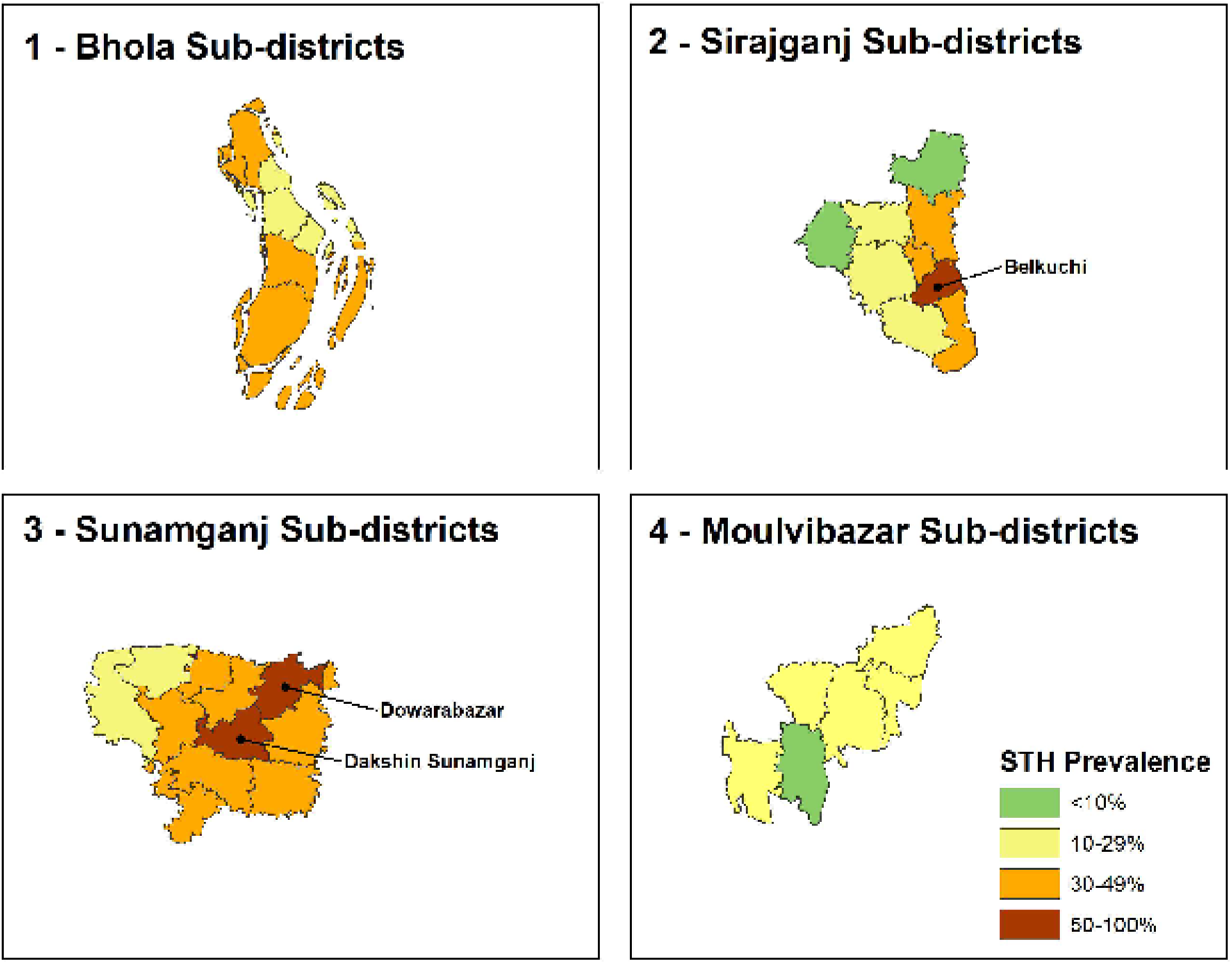
STH prevalence by sub-districts in the highest prevalent ICSPM districts.

### History of deworming

The proportion of self-reported deworming was highest among SAC (75.6%) followed by adults (69.1%) and PSAC (51.9%) for the 9,386 (86.7%) individuals who provided the history of deworming in the previous 6 months (table 2).

**Table 2:**
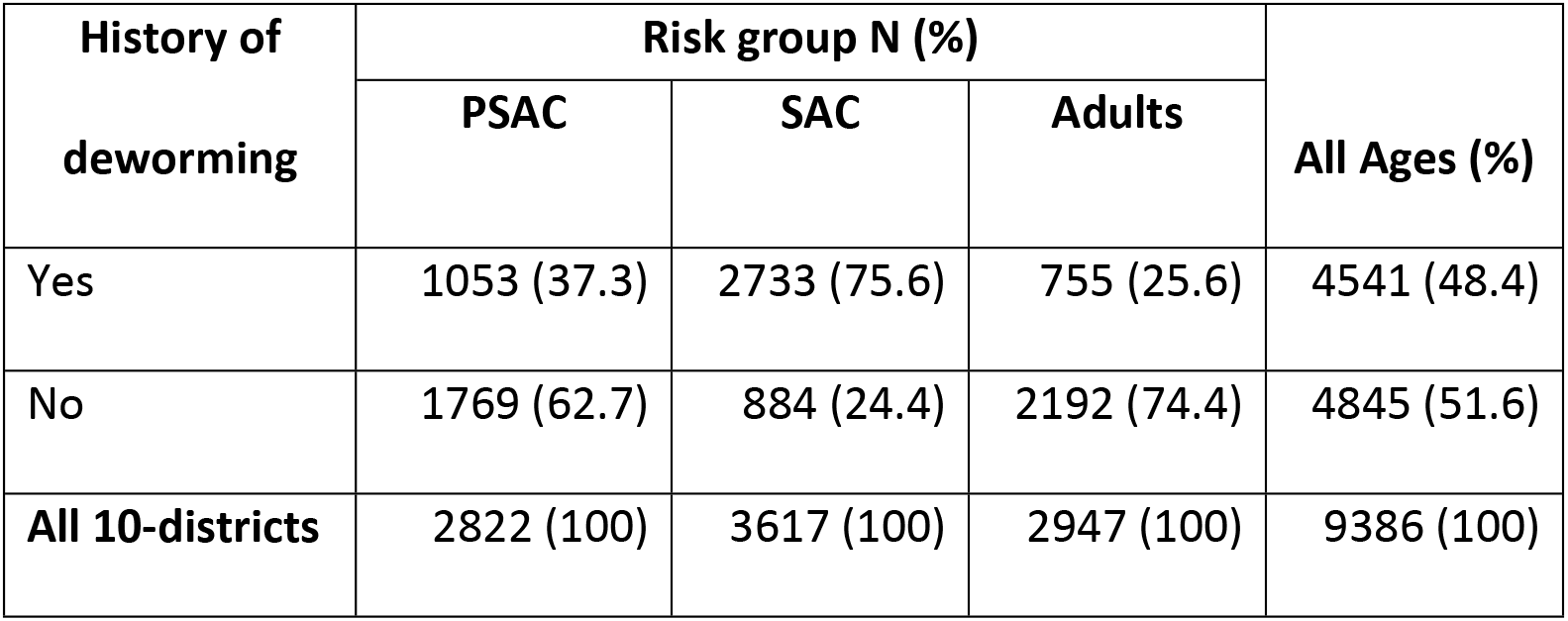
History of deworming within the past 6 months

Among responders (n= 7,469) to the query of the location of deworming, 88.6% of SAC reported getting dewormed through school-based MDA, while 85.1% adults and 76.2% of PSAC were dewormed through locally purchased deworming medicines (figure 5).

**Figure 5:**
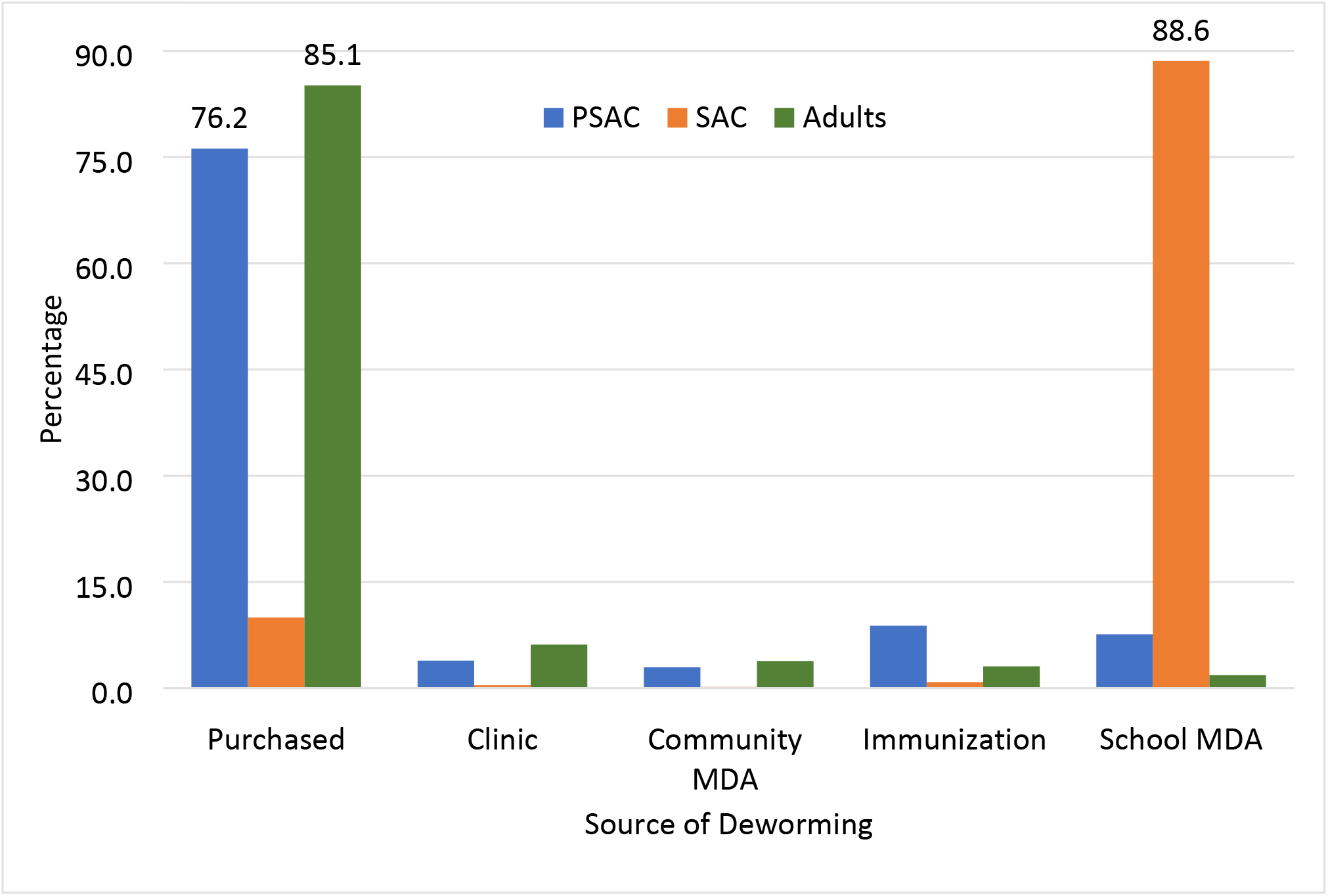
Source of deworming among those who reported deworming in the previous year.

## Discussion

To our knowledge, this is the first time an impact evaluation of MDA directed at one specific risk group, SAC, has shown a significant reduction in STH prevalence across all age groups in a given community. Our analysis of pooled data from community-based surveys in 10 districts in Bangladesh found a substantial reduction in overall STH prevalence from 79.8% (2005) to 14.0% (2017-2020) across all risk groups after more than 10 years of school-based systematic biannual PCT for SAC. Despite SAC being the only targeted risk group for MDA, the data shows no statistically significant differences in STH prevalence among PSAC, SAC, and adults. Although we did not specifically explore potential impact variables on the community prevalence, we speculate that the following factors might have contributed to this observation:

1. Change in health-seeking behavior in adults, namely purchasing deworming medication for themselves and family members outside of school (correlates with data on the source of deworming for PSAC and adults). This could be attributed to the positive results of school-based MDA encouraging out-of-school villagers to seek deworming, and
2. Improved WASH factors have increased access to improved sanitation at the household level.

In a recent national school survey of 5 to 12-year-old schoolchildren from 106 schools across the country (data not published yet), Bangladesh ELFSTH reported a lower STH prevalence than the prevalence derived from ICSPM surveys for the same age range (8.1% vs. 13%). Similarly, the MHII among this age group was 1.7% in the school-based survey and 2.6% with the ICSPM. The school-based survey had some limitations, including:

a. A small proportion of only school-going children were sampled as the targeted population rather than the entire endemic community.
b. The survey was not statistically powered to the program implementation level, nor the national school-attending population.

The observed difference in STH prevalence by school versus community-based surveys indicates that the school-based surveys may not be the right tool to estimate the true prevalence and intensity of STH for sub-national level program planning. This is especially true for advanced programs where prevalence is generally reduced and remaining aggregated areas of continuing transmission need to be reliably identified to redistribute precious resources for most efficient actions.

According to the latest WHO guidance(12), the 2030 goal for STH morbidity elimination is achieved when a country/region reaches <2% MHII. The school survey indicates that Bangladesh has achieved this goal and can halt MDA for 2 years per WHO guidance (13). However, the population-based ICSPM data for the same risk groups in the same geographic areas shows the true prevalence of intensity to be still > 2. The more granular ICSPM data provides more meaningful guidance to the program, i.e. reducing MDAs in low prevalence areas but increasing interventions in clusters of high transmission that persist. Our findings are similar to those from a school based survey in Sri Lanka (14).

Therefore, for countries with mature programs who have reach the WHO goal of consistent coverage above 75% for 5 years, we recommend a statistically valid, population-based sampling approach to assess the sub-national level impact on prevalence and intensity of STH for use in data-driven program decision-making or policy adjustments.

Additionally, our analysis revealed that the impact of STH control measures is not uniform across the country: it was significantly reduced in six districts, while the other four still carry a burden of higher prevalence and intensity. Potential factors influencing the impact of MDA on STH prevalence and intensity may be related to the local population and individual characteristics, as well as service processes related to intervention quality such as:

1. Varied baseline STH prevalence and intensity. (data not available)
2. Population movement across district borders, bringing the infection from other areas.
3. The complicated relationship between drug distributors and targeted risk groups.
4. Variable environmental or ecological characteristics among districts that support longer survival of STH eggs in the soil.
5. False rumors or distrust of government programs about the ‘real’ purpose of the treatment, and
6. Responses to local socio-cultural control measures.

Among high STH prevalent districts, STH prevalence ranged from 5.1% (Kazipur sub-district, Sirajganj) to 71% (Dowara Bazar sub-district, Sunamganj). All corresponding sub-districts (Sunamganj and Bhola), seven of nine sub-districts (Sirajganj), and one of seven sub-districts (Moulvibazar) had a prevalence of more than 20%.

The Bangladesh national NTD program’s STH control office plans to use these findings to design a focused intervention program in select sub-districts to lower unexpected high prevalence. Additionally, the national program may use these findings to make decisions for altering the frequency of MDA programs in low prevalence districts.

To further explore the consistent impact across all age groups, we reviewed the pill intake and their source data by age group and by district as well as collectively for all 10 districts. This revealed that a large proportion of PSAC and adults reported the purchase of locally available drugs as the primary source of deworming in the past 6 months. This finding has potential implications for the national program as it may indicate communal behavioral change towards self-investment in preventive health. This could be a spillover effect of the sustained impact of school deworming in these districts, signaling that school-based MDA and accompanying community messaging raises awareness of the positive health outcomes of deworming, triggering treatment-seeking behavior in community members who do not have access to school MDAs but do have local access to affordable, high quality deworming medicines (Bangladesh generic manufacturers and formulations of benzimidazoles: CWW web-survey, 2019). These initial findings need further exploration and, if confirmed, will be an important factor influencing a national policy of sustainable domestic financing guided by quality disease, socio-behavioral and pharmacological data. Similar behavioral changes should be explored by other national programs that have quality generics available locally for deworming.

## Conclusion

After 23 rounds of school-based MDA to lower the burden of STH infection since 2008, a review of survey data from 10 districts in Bangladesh shows that it is close to eliminating the infection as a public health problem from most of the country. To sustain current progress and move forward, Bangladesh needs to identify and treat all community members at risk in the persistent high-prevalence pockets of geographic areas, such as Sunamganj, Bhola, and Sirajganj. Community-based surveys may serve as better tools for a true assessment of PCT in endemic communities compared to more common school-based surveys and additional information on deworming sources is a valuable resource for national programs with high, multi-year coverage.

## Limitations

The ICSPM surveys had some limitations including a higher than expected stool nonresponse rate, possible recall bias (particularly the responses to the history and location of deworming questions), and gender inequity among adult respondents. Additionally, the timing between the stool sample deposit by the survey respondents and testing in the laboratory may have been longer than “ideal” due to geographical challenges so may have underestimated the hookworm prevalence slightly but there is only one published study that documents the “ideal” specimen testing interval for hookworms (15) and additional studies have shown little or no hookworm in south Asia.

It is of note that Bangladesh’s ELFSTH treated 19 LF-endemic districts with Albendazole (also active against STH worms) to control Lymphatic Filariasis (LF) through community-based MDAs from 2001 to 2014. While these LF-focused MDAs also impacted the STH prevalence in those 19 districts, these treatments did not affect ICSPM results as the LF program ceased in 2014 and ICSPM started data collection in 2017.

## Acknowledgments

We would like to express our sincere gratitude to all the staff of the respective district & sub-district health officials, field staff, and survey participants in Bangladesh, as well as the multiple CWW team members and consultants who historically contributed to protocol development and early implementation of some surveys. These surveys were supported with generous funding from Johnson & Johnson, and the Government of Canada.

